# Beta human papillomavirus 8 E6 allows colocalization of non-homologous end joining and homologous recombination repair factors

**DOI:** 10.1101/2021.06.11.448030

**Authors:** Changkun Hu, Taylor Bugbee, Rachel Palinski, Nicholas Wallace

## Abstract

Beta human papillomavirus (β-HPV) are hypothesized to make DNA damage more mutagenic and potentially more carcinogenic. Double strand breaks in DNA (DSBs) are the most deleterious DNA lesion. They are typically repaired by homologous recombination (HR) or non-homologous end joining (NHEJ). HR occurs after DNA replication while NHEJ can occur at any point in the cell cycle. They are not thought to occur in the same cell at the same time. By destabilizing p300, β-HPV type 8 protein E6 (β-HPV8 E6) attenuates both repair pathways. However, β-HPV8 E6 delays rather than abrogates DSB repair. Thus, β-HPV8 E6 may cause DSBs to be repaired through a more mutagenic process. To evaluate this, immunofluorescence microscopy was used to detect colocalization, formation, and resolution of DSB repair complexes following damage. Flow cytometry and immunofluorescence microscopy were used to determine the cell cycle distribution of repair complexes. The resulting data show that β-HPV8 E6 causes HR factors (RPA70 and RAD51) to colocalize with a persistent NHEJ factor (pDNA-PKcs). RPA70 complexes gave way to RAD51 complexes as in canonical HR, but RAD51 and pDNA-PKcs colocalization did not resolve within 32 hours of damage. The persistent RAD51 foci occur in G1 phase, consistent with recruitment after NHEJ fails. Chemical inhibition of p300, p300 knockout cells, and an β-HPV8 E6 mutant demonstrate that these phenotypes are the result of β-HPV8 E6-meidated p300 destabilization. Mutations associated with DSB repair were identified using next generation sequencing after a CAS9-induced DSB. β-HPV8 E6 increases the frequency of mutations (>15 fold) and deletions (>20 fold) associated with DSB repair. These data suggest that β-HPV8 E6 causes abnormal DSB repair where both NHEJ and HR occur at the same lesion and that his leads to deletions as the single stranded DNA produced during HR is removed by NHEJ.

**Author Summary:** Our previous work shows that a master transcription regulator, p300, is required for two major DNA double strand break (DSB) repair pathways: non-homologous end joining (NHEJ) and homologous recombination (HR). By degrading p300, beta genus Human Papillomavirus 8 protein E6 (β-HPV8 E6) hinders DNA-PKcs activity, which is a key factor of NHEJ. β-HPV8 E6 also stalls HR via p300 degradation resulting in the persistence of a core factor, RAD51. NHEJ and HR are known competitive to each other and only one pathway can be initiated to repair a DSB. Particularly, NHEJ tends to be used in G1 phase and HR occurs in S/G2 phase. Here, we show that β-HPV8 E6 allows NHEJ and HR to occur at the same break site. This is expected to be mutagenic because HR generates overhangs while NHEJ removes them. Further, we show that β-HPV8 E6 allows HR to occur in G1, which cannot be finished due to the lack of homologous templates. Finally, our sequencing results show that β-HPV8 E6 significantly increases genomic variations including deletions and insertions following CAS9 induced DSB. This study supports the hypothesis that β-HPV8 infections increases genomic instability.

## Introduction

Beta genus of human papillomavirus (β-HPVs) are frequently found in human skin [1,2]. HPV replication requires actively proliferating cells and the replication machinery of the host cells, which puts β-HPV infections in conflict with the cell cycle arrest associated with the repair of UV photolesions [3–6]. Potentially as a mechanism to counter cell cycle arrest, some β-HPVs hinder the cellular response to DNA damage [7–9]. The E6 gene from β-HPV type 8 (β-HPV8 E6) dysregulates the cellular response to DNA damage by binding and destabilizing p300, a histone acetyltransferase that regulate transcription by chromosome remodelling [10–12]. p300 destabilization decreases expression of at least four DNA repair genes (ATM, ATR, BRCA1, and BRCA2) [7,13,14]. β-HPV8 E6 also reduces ATM and ATR activation in response to UV [8]. This hinders UV damage repair, making UV-induced double stranded DNA breaks (DSBs) more likely [14].

DSBs are the most deleterious type of DNA lesion. Among DSB repair pathways, non-homologous end joining (NHEJ) and homologous recombination (HR) are best characterized and associated with cell cycle arrest. NHEJ can happen throughout the cell cycle but tends to occur during G1 and early S phase [15–18]. NHEJ initiation occurs when a DSB is sensed and Ku70 and Ku80 bind at the lesion [16,19]. DNA-dependent protein kinase catalytic subunit (DNA-PKcs) is then recruited to form a heterotrimer known as the DNA-PK holoenzyme. This allows activation of DNA-PKcs via autophosphorylation at S2056 (pDNA-PKcs) and facilitates downstream steps in the pathway, including Artemis activation [20–22]. Artemis has both endonuclease and exonuclease activity that remove single stranded DNA, producing blunt ends [23–25]. Once processed, other NHEJ factors (e.g. XRCC4, XLF, and DNA ligase IV) ligate the gap to resolve the DSB [26–28].

Homologous recombination uses a sister chromatid as a homologous template to provide error-free DSB repair, typically restricting the pathway to the S and G2 phases [29–32]. HR initiation includes the formation of a Mre11, RAD50, and NBS1 heterotrimer, known as the MRN complex [33]. The MRN complex resects DNA around the DSB, resulting in single stranded DNA overhangs [33–35]. An RPA trimer (RPA70, RPA32, and RPA14) rapidly coats this single stranded DNA, helping maintain the stability of the DNA [30,36–38]. Then BRCA1, BRCA2, and PALB2 facilitate the exchange of RPA trimers for RAD51 [39]. Once loaded onto DNA near a DSB, RAD51 facilitates a search for the homologous template, strand invasion, and helps resolve the lesion [31,40,41].

β-HPV8 E6 attenuates the overall efficiency of both NHEJ and HR [7,9]. While β-HPV8 E6 does not cause any known defects in NHEJ or HR initiation, it prevents resolution of pDNA-PKcs and RAD51 repair complexes by destabilizing p300 [7,9]. These observations are consistent with the hypothesized ability of some β-HPV infections to promote skin cancer, by making UV damage more mutagenic [42,43]. However, given the transient nature of β-HPV infections, there remains uncertainty regarding whether β-HPV infections are sufficiently genotoxic to introduce enough mutations to drive tumorigenesis after the viral infection is cleared.

Here, we use immunoblotting, immunofluorescence microscopy, and flow cytometry to better characterize the persistent pDNA-PKcs and RAD51 repair complexes previously reported in cells expressing β-HPV8 E6 [7,9]. NHEJ and HR are mechanistically incompatible; and therefore, NHEJ (pDNA-PKcs) and HR (RAD51) typically do not occur at the same break site at the same time [44]. However, we show that β-HPV8 E6 promotes the colocalization of pDNA-PKcs and RAD51 foci. This abnormal colocalization is caused by the destabilization of p300 and occurs more frequently in cells with more β-HPV8 E6. Using p300 knockout cell lines, a p300 inhibitor, and a mutated β-HPV8 E6, we show that loss of p300 allows HR to initiate in G1 and at sites of failed NHEJ. Finally, we developed an assay that combines sgRNA/CAS9 endonuclease activity and next generation sequencing to define the extent to which β-HPV8 E6 increased mutations during DSB repair. This approach demonstrated β-HPV8 E6 caused a greater than ten-fold increase in mutations in the 200 Kb surrounding a DSB.

## Results

### β-HPV8 E6 promotes the recruitment of HR factors to sites of stalled NHEJ

DSBs are typically repaired in human foreskin keratinocytes within two to eight hours after they occur [7]. However, β-HPV8 E6 causes DSBs to persist for at least twenty-four hours by delaying the resolution of HR (RAD51) and NHEJ (pDNA-PKcs) repair complexes [7,9]. These studies demonstrate that β-HPV8 E6 attenuated HR and NHEJ repair. However, they did not characterize the persistent repair complexes. Vector control (HFK LXSN) or β-HPV8 E6 (HFK β-HPV8 E6) expressing telomerase immortalized human foreskin keratinocyte cell lines were used to address this knowledge gap [45]. β-HPV8 E6 was HA-tagged and the expression was confirmed using immunofluorescence microscopy (S1A Fig). DSBs were induced in these cells by growth in media containing 10 µg/mL of zeocin for 10 min, a water-soluble radiation mimetic [46]. Twenty-four hours after removal from zeocin-containing media, immunofluorescence microscopy was used to detect RAD51 and pDNA-PKcs repair complexes. Consistent with numerous reports that NHEJ and HR are employed at separate phases of the cell cycle [47–49], HFK LXSN cells infrequently contained both RAD51 and pDNA-PKcs foci (S1B-C Fig). However, HFK β-HPV8 E6 cells were significantly more likely to have both RAD51 and pDNA-PKcs foci in the same cells (S1B-C Fig). Moreover, β-HPV8 E6 significantly increased the colocalization of RAD51 and pDNA-PKcs repair complexes (Fig 1A-B).

**Fig 1.**
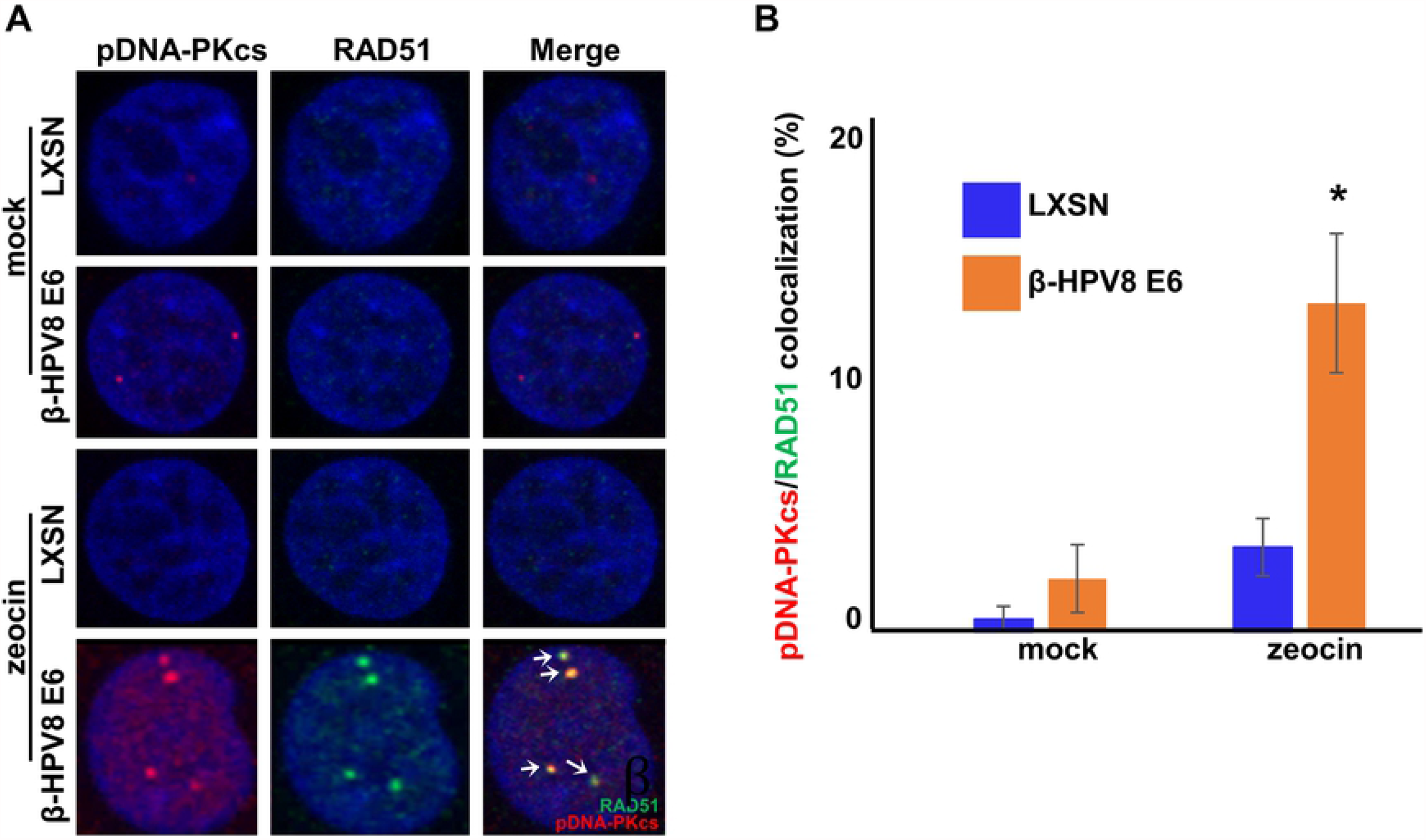
β-HPV8 E6 allows RAD51/pDNA-PKcs colocalization. (A) Representative image of HFK LXSN and HFK β-HPV8 E6 cells stained for RAD51 (green) and pDNA-PKcs (red) 24 hours after DSB induction by growth in media containing zeocin (10 µg/mL, 10 min) or media containing additional water (solvent for zeocin) as a negative control. Additional water control is described as “mock” in Fig. White arrows indicate colocalizing RAD51 and pDNA-PKcs foci. (B) Percentage of HFK cells with colocalized RAD51 and pDNA-PKcs foci after mock treatment or zeocin treatment. All values are represented as mean ± standard error from three independent experiments. The statistical significance of differences between cell lines were determined using Student’s t-test. * indicates p-value < 0.05 (n=3). All microscopy images are 400X magnification.

RAD51 and pDNA-PKcs foci colocalization is rarely reported and has not been thoroughly characterized [50,51]. pDNA-PKcs repair foci represent an intermediate step in the NHEJ pathway, while RAD51 repair complex formation represents a near terminal step in the HR pathway. RAD51 foci appear after the formation and resolution of RPA70 repair complexes [52]. This suggested that RPA70 repair complexes also colocalized with pDNA-PKcs foci. Immunofluorescence microscopy was used to test this. HFK LXSN and HFK β-HPV8 E6 cells were stained for RPA70 and pDNA-PKcs at intervals over a 32-hour period after cells were removed from zeocin containing media. RPA70 and pDNA-PKcs rarely colocalized in HFK LXSN cells but their colocalization was readily detectable in HFK β-HPV8 E6 cells (Fig 2A-B). The peak of RPA70/pDNA-PKcs foci colocalization in HFK β-HPV8 E6 cells occurred 16-hours after DSB induction. RAD51/pDNA-PKcs colocalization was similarly detected over a 32-hour period after zeocin treatment (Fig 2C-D). Consistent with active progression through the HR pathway, the peak for RAD51/pDNA-PKcs colocalization occurred 24-hours after cells were removed from zeocin containing media. The cell line specificity of these observations was probed by examining DSB repair in empty vector control and β-HPV8 E6 expressing osteosarcoma cells, U2OS LXSN and U2OS β-HPV8 E6, respectively. β-HPV8 E6 expression was confirmed in these cells by probing for p300. As previously reported [9], U2OS β-HPV8 E6 cells had less p300 than U2OS LXSN cells (S2A Fig.). β-HPV8 E6 maintained its ability to promote RPA70 and RAD51 colocalization with pDNA-PKcs in U2OS cells (Fig 2E-F and S2B-C Fig.). Further, the kinetics of RPA70 and RAD51 recruitment to pDNA-PKcs foci were similar in HFK β-HPV8 E6 and U2OS β-HPV8 E6 cells.

**Fig 2.**
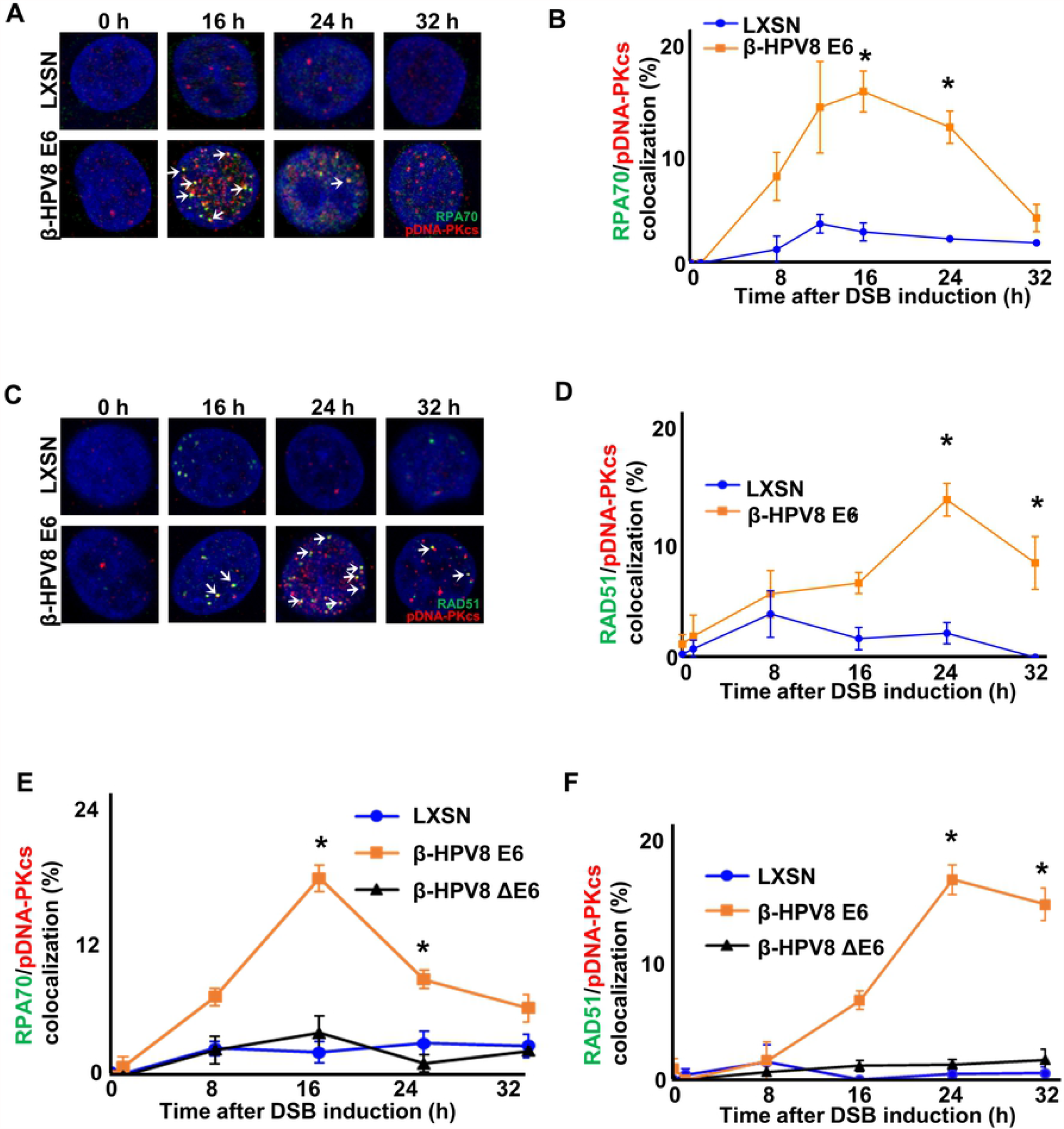
By binding p300, β-HPV8 E6 allows HR foci to colocalize with persistent pDNA-PKcs foci. (A) Representative images of HFK LXSN and HFK β-HPV8 E6 cells stained for RPA70 (green) and pDNA-PKcs (red) following zeocin treatment (10 µg/mL, 10 min). (B) Percentage of HFK cells with colocalized RPA70 and pDNA-PKcs foci. (C) Representative images of HFK LXSN and HFK β-HPV8 E6 cells stained for RAD51 (green) and pDNA-PKcs (red) following zeocin treatment (D) Percentage of HFK cells with colocalized RAD51 and pDNA-PKcs foci. (E) Percentage of U2OS cells with colocalized RPA70 and pDNA-PKcs foci following zeocin treatment (100 µg/mL, 10 min). (F) Percentage of U2OS cells with colocalized RAD51 and pDNA-PKcs foci following zeocin treatment. White arrow indicates colocalization. All values are represented as mean ± standard error from three independent experiments. The statistical significance of differences between LXSN and β-HPV8 E6 cell lines were determined using Student’s t-test. * indicates p < 0.05 (n=3). All microscopy images are 400X magnification.

### Catalytic activity of p300 prevents colocalization of HR factors with persistent pDNA-PKcs repair complexes

β-HPV8 E6 obtains most of its known DNA repair inhibitory effects by binding p300. This includes its ability to impair the HR and NHEJ pathways [7,9]. Thus, a mutant β-HPV8 E6 (β-HPV8 ΔE6) that has the p300 binding site (residues 132-136) deleted has significantly less ability to restrict DNA repair [10,14]. To determine if β-HPV8 E6 promoted the colocalization of HR factors with pDNA-PKcs by destabilizing p300, β-HPV8 ΔE6 was expressed in U2OS cells (U2OS β-HPV8 ΔE6) [7]. The expression of β-HPV8 ΔE6 did not result in reduced p300 abundance (S2A Fig.). Consistent with a p300-dependent phenotype, colocalization of RPA70 and RAD51 with pDNA-PKcs was rarely detected in U2OS β-HPV8 ΔE6 by immunofluorescence microscopy after zeocin-induced DSBs (Fig 2E-F and S2BC Fig.). Indeed, the colocalization frequency was indistinguishable in U2OS β-HPV8 ΔE6 and U2OS LXSN cells (Fig 2E-F, and S2B-C Fig.).

Because deletion of the p300 binding domain has been reported to disrupts other functions of β-HPV8 E6 [53], HCT116 cells with wild type p300 (HCT116 P300 WT) and HCT116 cells with the p300 gene knocked out (HCT116 KO) were used to verify the role of p300 in preventing the colocalization of NHEJ and HR repair factors [12]. Immunoblotting was used to confirm p300 knockout (S3A Fig.). DSBs were induced by zeocin (10 µg/mL for 10 min). Immunofluorescence microscopy was used to detect colocalization of RPA70 and pDNA-PKcs over a 32-hour period after zeocin treatment. RPA70/pDNA-PKcs colocalization peaked 16-hours after DSB induction in both cell lines but was significantly higher in HCT116 P300 KO cells (Fig 3A and 3B). The frequency of RAD51/pDNA-PKcs colocalization was also significantly higher in HCT116 P300 KO cells than HCT116 P300 WT cells (Fig 3C and 3D). RAD51/pDNA-PKcs colocalization peaked after the RPA70/pDNA-PKcs colocalization peaks, indicating the progression through the HR pathway.

**Fig 3.**
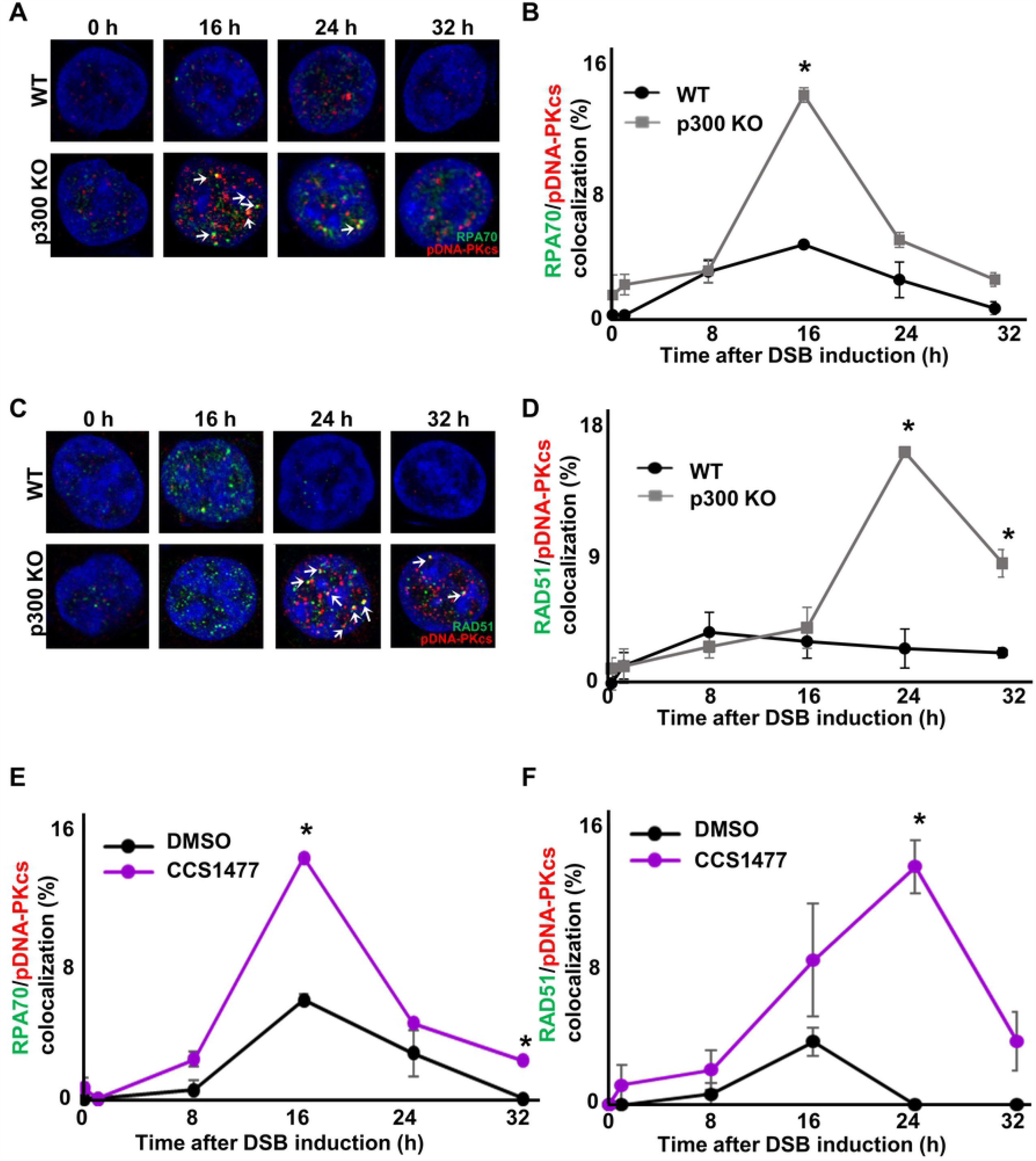
p300 reduces co-localization of HR foci with pDNA-PKcs foci. (A) Representative images of HCT116 p300 WT and HCT116 p300 KO cells stained for RPA70 (green) and pDNA-PKcs (red) following zeocin treatment (100 µg/mL, 10 min). (B) Percentage of HCT116 cells with colocalized RPA70 and pDNA-PKcs foci. (C) Representative images of HCT116 p300 WT and HCT116 p300 KO cells stained for RAD51 (green) and pDNA-PKcs (red) following zeocin treatment. (D) Percentage of HCT116 cells with colocalized RAD51 and pDNA-PKcs foci. (E) Percentage of HFK LXSN cells treated with CCS1477 (1 μM) or DMSO that contained colocalized RPA70 and pDNA-PKcs foci following zeocin treatment (10 µg/mL, 10 min). (F) Percentage of HFK LXSN cells treated with CCS1477 (1 μM) or DMSO that contained colocalized RAD51 and pDNA-PKcs foci following zeocin treatment (10 µg/mL, 10 min). White arrow indicates colocalization. All values are represented as mean ± standard error from three independent experiments. The statistical significance of differences between cell lines or treatments were determined using Student’s t-test. * indicates p < 0.05 (n=3). All microscopy images are 400X magnification.

p300 is a large (∼300kDa) protein that can promote repair by acting as a scaffold for other repair factors or through its catalytic activity [54]. To determine the extent that the catalytic activity of p300 restricted the colocalization of pDNA-PKcs with RAD51 and RPA70, HFK LXSN cells were grown in media containing a small molecule p300 inhibitor (1 µM of CCS1477). A previous study showed that p300 is required for ATM and ATR activation [8]. As a confirmation of p300 inhibition, CCS1477 restricted damage-induced ATM and ATR phosphorylation (S4A Fig.). CCS1477 also significantly increased RPA70/pDNA-PKcs and RAD51/pDNA-PKcs colocalization after zeocin-induced DSBs (Fig 3E-F, S5A-B Fig.). Again, colocalization of RPA70/pDNA-PKcs peaked before colocalization of RAD51/pDNA-PKcs. Together, these data show that p300 activity reduces the frequency with which HR factors are recruited to sites of persistent pDNA-PKcs foci.

### β-HPV8 E6 allows RAD51 foci formation in G1 phase by binding to p300

These data suggest that the colocalization of HR factors with pDNA-PKcs occurs as a result of cells initiating and then failing to complete NHEJ. Because NHEJ is the dominant pathway during G1 phase, the failed NHEJ caused by β-HPV8 E6 likely occurs during G1. Thus, β-HPV8 E6 may be promoting the formation of RAD51 foci during G1 when HR is unlikely to be completed due to the absence of sister chromatids. Immunofluorescence microscopy was used to evaluate this possibility, with cyclin E serving as a marker of cells in G1 [55,56]. Twenty-four hours after DSB induction, co-staining of cyclin E and RAD51 foci were significantly more frequent in HFK β-HPV8 E6 than in HFK LXSN cells (Fig 4A-B). Next, flow cytometry was used to define the frequency of RAD51 positive cells in G1. DAPI staining was used to define relative DNA content and identify cells in G1. Antibodies against RAD51 (along with the appropriate secondary antibody) were used to label cells with increased RAD51. Cells were stained with only secondary antibody to determine the background level of RAD51 staining (S6A Fig.). After zeocin-induced DSBs (0, 4 and 24 hours later), the fraction of cells in G1 was analysed for RAD51 staining. As a confirmation that RAD51 staining was damage-induced, growth in zeocin-containing media significantly increased the frequency of RAD51 staining in HFK LXSN (S6B Fig.). Consistent with prior reports [57,58], RAD51 staining was rarely detected in HFK LXSN cells during the G1 (Fig 4C and 4D). However, there was a significant increase in HFK β-HPV8 E6 cells that stained positive for RAD51 during G1. To confirm that these results were not cell type specific, the frequency of RAD51 staining in G1 was similarly probed in U2OS cells. U2OS β-HPV8 E6 cells were more likely to have RAD51 staining during G1 than U2OS LXSN (Fig 4E-F and S7A-C Fig.).

**Fig 4.**
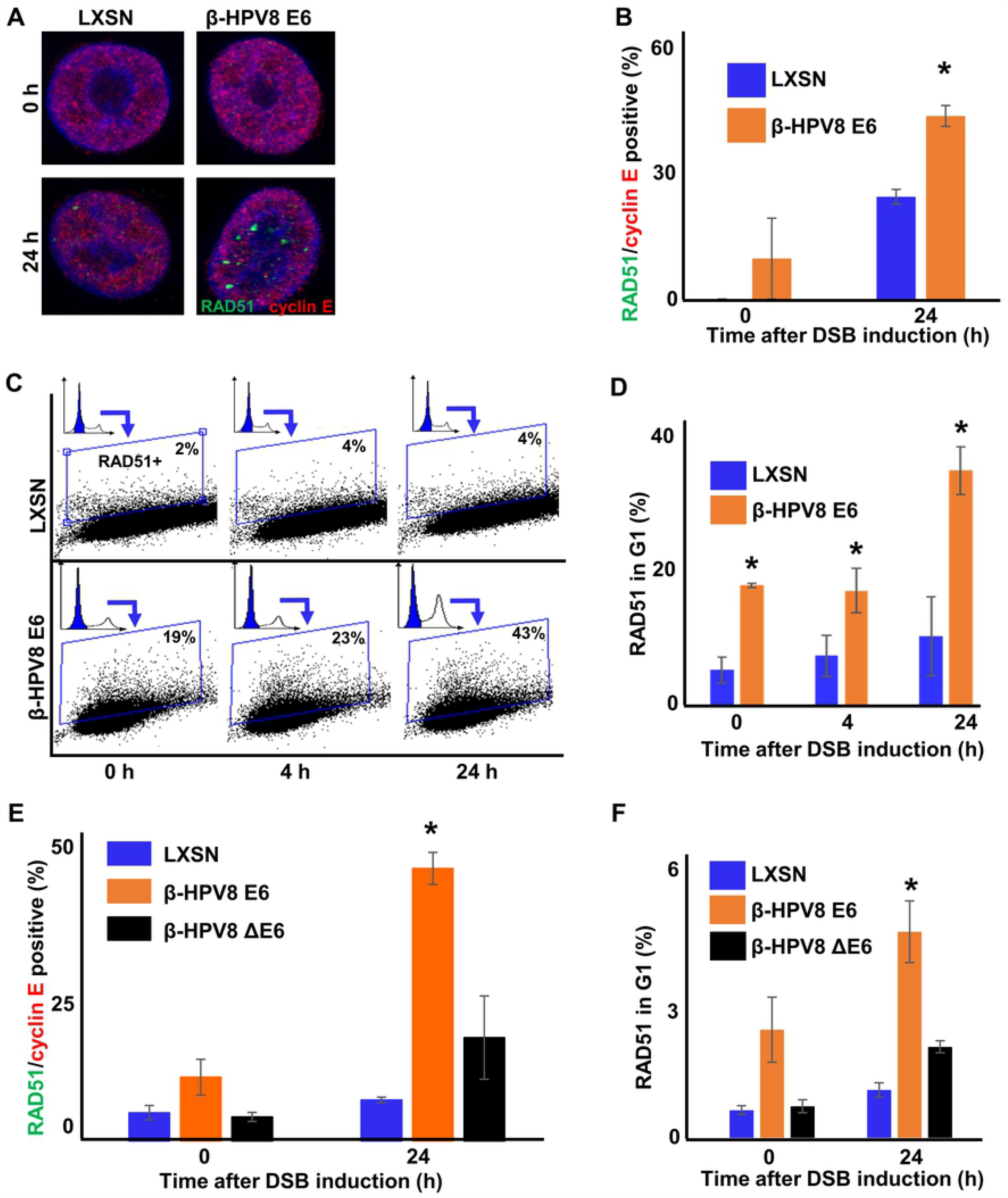
β-HPV8 E6 allows RAD51 foci formation in G1. (A) Representative cyclin E positive HFK LXSN and HFK β-HPV8 E6 cells stained for RAD51 (green) and cyclin E (red) 24 hours following zeocin treatment (10 µg/mL, 10 min). (B) Percentage of cyclin E positive HFK cells with RAD51 foci after zeocin treatment. (C) Representative images of flow cytometry analysis of HFK LXSN and HFK β-HPV8 E6 cells stained with DAPI and RAD51. DAPI was used to identify G1 phase (blue area in inset) that were then gated on RAD51 intensity. The gating represents RAD51 positive based off secondary only control. (D) Percentage of HFK cells in G1 that are positive for RAD51. (E) Percentage of cyclin E positive U2OS cells with RAD51 foci after zeocin treatment (100 µg/mL, 10 min). (F) Percentage of U2OS cells in G1 that are positive for RAD51 by flow cytometry. All values are represented as mean ± standard error from three independent experiments. The statistical significance of differences between LXSN and β-HPV8 E6 cell lines were determined using Student’s t-test. * indicates p < 0.05(n=3). All microscopy images are 400X magnification.

Consistent with a p300-dependent mechanism of action, RAD51 staining was less common in U2OS β-HPV8 ΔE6 in G1 compared to U2OS β-HPV8 E6 (Fig 4E-F and S7A-C Fig). HCT116 P300 WT and HCT116 P300 KO cells were used to more rigorously evaluate the ability of p300 to prevent RAD51 foci formation during G1. These cells were stained for RAD51 and cyclin E 24-hours after DSB induction and visualized by immunofluorescence microscopy. Co-staining of cyclin E and RAD51 was significantly more likely to occur in HCT116 P300 KO than HCT116 p300 WT cells (Fig 5A and 5B). Further, flow cytometry analysis was used to determine the frequency of RAD51 repair staining in G1. As described before, cells were stained with only secondary antibody to determine the background level of RAD51 staining (S8A Fig). As a confirmation that RAD51 staining was damage-induced, growth in zeocin-containing media significantly increased the frequency of RAD51 staining in HFK LXSN (S8B Fig). HCT116 p300 KO cells were significantly more likely to be RAD51 positive during G1 than HCT116 p300 WT cells (Fig 5C-D). To examine the role of p300 activity in preventing RAD51 foci formation during G1, HFK LXSN cells were grown in media containing CCS1477 (1 µM). CCS1477 increased the frequency with which HFK LXSN cells contained RAD51 foci and stained positive for cyclin E (Fig 5E and S9A Fig.). Flow cytometry analysis also showed that CCS1477 significantly increased the frequency of RAD51 positive cells in G1 phase (Fig 5F and S9B-C Fig.). Together these data demonstrate that p300 activity prevents RAD51 foci formation during G1 and that β-HPV8 E6 can abrogate this protection by binding p300.

**Fig 5.**
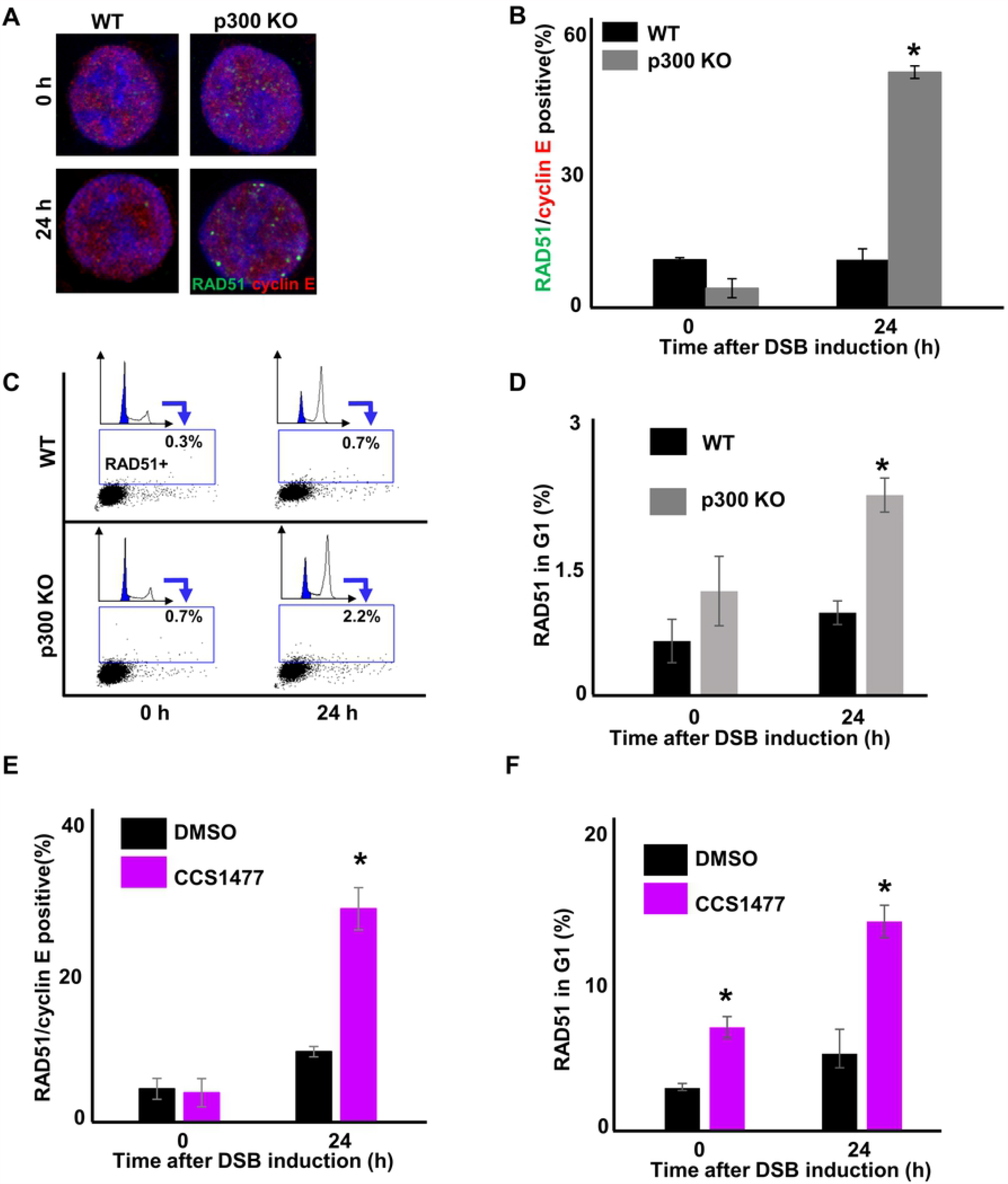
p300 restricts RAD51 foci formation in G1. (A) Representative cyclin E positive HCT116 p300 WT and HCT116 p300 KO cells stained for RAD51 (green) and cyclin E (red) following zeocin treatment (100 µg/mL, 10 min). One cyclin E negative and one cyclin E positive cell are showed for each cell line. (B) Percentage of cyclin E positive HCT116 cells with RAD51 foci following zeocin treatment. (C) Representative images of flow cytometry analysis in HCT116 p300 WT and HCT116 p300 KO cells stained for DAPI and RAD51 following zeocin treatment. DAPI was used to identify G1 populations (blue area in inset) that were then gated on RAD51 intensity. The gating represents RAD51 positive based off secondary only control. (D) Percentage of HCT116 cells in G1 that are positive for RAD51. (E) Percentage of cyclin E positive HFK cells treated with DMSO or CCS1477 (1 μM) positive for RAD51 foci following zeocin treatment (10 µg/mL, 10 min). (F) Percentage of HFK cells treated with DMSO or CCS1477 in G1 that are positive for RAD51 following zeocin treatment (10 µg/mL, 10 min). All values are represented as mean ± standard error from three independent experiments. The statistical significance of differences between cell lines or treatments were determined using Student’s t-test. * indicates p < 0.05 (n=3). All microscopy images are 400X magnification.

### DNA-PKcs inhibition leads to HR initiation in G1 phase

These data above suggest that impaired NHEJ leads to HR initiation during G1. To define the extent that stalled NHEJ resulted in persistent RAD51 foci, HFK LXSN cells were grown in media containing a well-characterized DNA-PKcs inhibitor ((1 µM of NU7441), S10A Fig.). As a control, immunofluorescence microscopy was used to detect a standard marker of DSBs (phosphorylation of H2AX at serine 139 or pH2AX) at 0-, 1-, and 24-hour time points after DSB-induction. Compared to cells grown in media containing DMSO (solvent), NU7441 delayed pH2AX foci resolution (S10B-C Fig). Next, the extent to which NU7441 changed the kinetics of RAD51 foci resolution was determined. HFK LXSN cells were again grown in media containing NU7441 or DMSO and stained 0-, 1-, and 24-hours after the induction of DSBs. Immunofluorescence microscopy was used to detect RAD51 foci and demonstrated that NU7441 increased the persistence of RAD51 foci (Fig 6A-B). This is consistent with impaired NHEJ leading to stalled HR.

**Fig 6.**
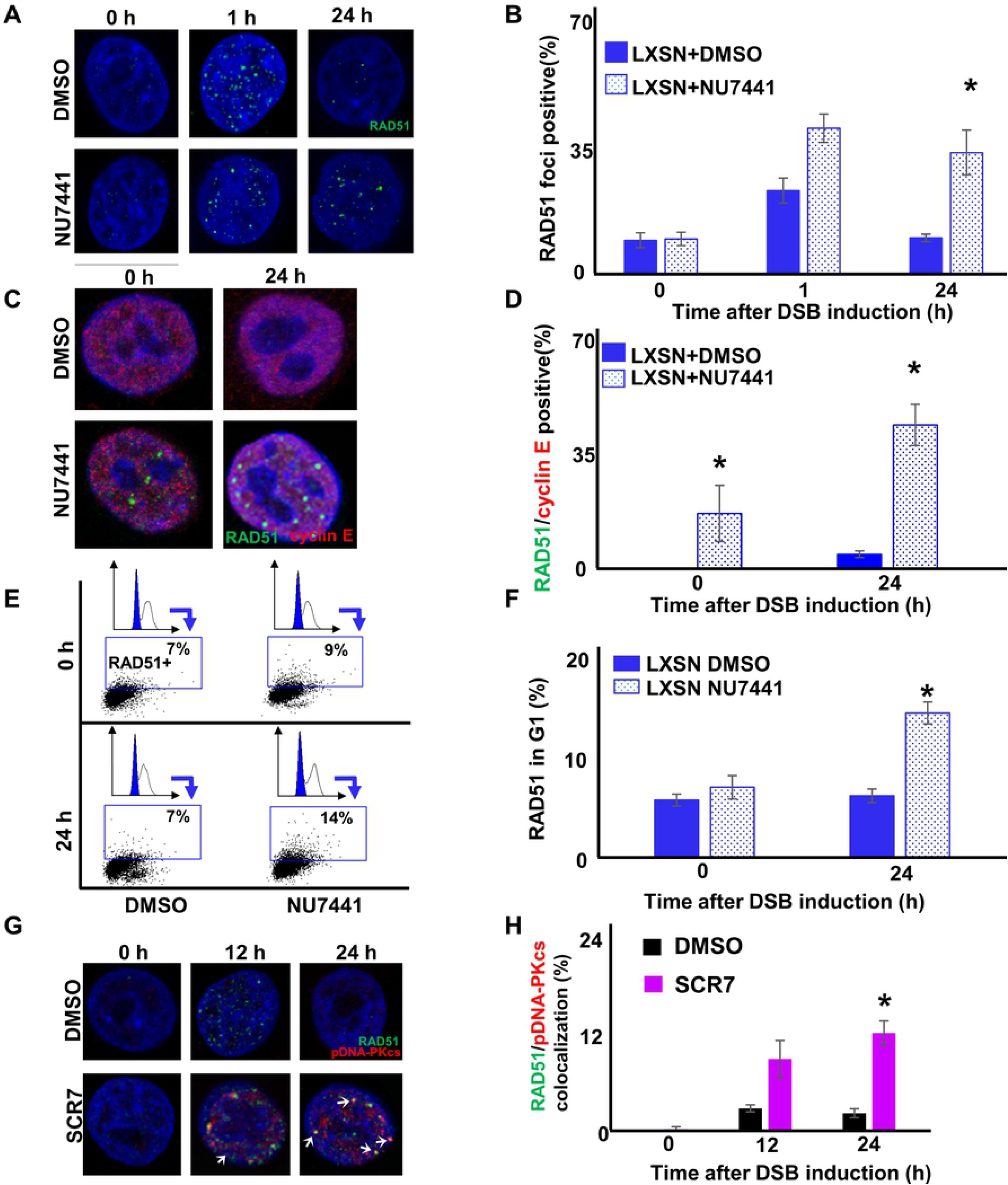
DNA-PKcs inhibition increases RAD51 foci in G1. (A) Representative images of HFK LXSN cells treated with NU7441 (1 µM) or DMSO stained for RAD5 (green) following zeocin treatment (10 µg/mL, 10 min). (B) Percentage of HFK LXSN cells with RAD51 foci at indicated times following zeocin treatment. (C) Representative cyclin E positive HFK LXSN cells treated with NU7441 or DMSO stained for cyclin E and RAD51 following zeocin treatment. (D) Percentage of HFK LXSN cells positive for cyclin E and RAD51 foci following zeocin treatment (E) Representative images of flow cytometry analysis in HFK LXSN cells treated with NU7441 following zeocin treatment. DAPI was used to identify G1 populations (blue area in inset) that were then gated by RAD51 intensity. The gating represents RAD51 positive based off secondary only control. (F) Percentage of LXSN HFK cells in G1 that are positive for RAD51 following zeocin treatment. (G) Representative images of HFK LXSN cells treated with ligase IV inhibitor (1 µM of SCR7) or DMSO stained for RAD51 and pDNA-PKcs at indicated time points following zeocin treatment. (H). Percentage of HFK LXSN cells with colocalized RAD51 and pDNA-PKcs foci at indicated time points after treatment with ligase IV inhibitor or DMSO and exposure to zeocin. White arrow indicates colocalization. All values are represented as mean ± standard error from three independent experiments. The statistical significance of differences between treatments were determined using Student’s t-test. * indicates p < 0.05 (n=3). All microscopy images are 400X magnification.

To test the hypothesis that impaired NHEJ leads to HR initiation during G1, cyclin E and RAD51 foci were detected by immunofluorescence microscopy with and without NU7441 (1 µM). Twenty-four hours after zeocin treatment, NU7441 increased the frequency of HFK LXSN cells that contained RAD51 foci and stained positive for cyclin E (Fig 6C-D). Moreover, flow cytometry analysis was used to detect cell stained positive for RAD51 G1. As described above, cells stained with only secondary antibody to determine the background level of RAD51 staining (S11A Fig.). As a confirmation that RAD51 staining was damage-induced, zeocin treatment significantly increased the frequency of RAD51 staining in HFK LXSN (S11B Fig.). NU7441 significantly increased RAD51 staining in G1 (Fig 6E-F). To test cell line specificity, NU7441 made co-staining for RAD51 foci and cyclin E more frequent in U2OS LXSN (S12A Fig.) and HCT116 WT cells (S12B Fig.). These data demonstrate that DNA-PKcs inhibition results in abnormal RAD51 repair complex formation during G1. However, because NU7441 prevents phosphorylation of DNA-PKcs. Thus, colocalization experiments could not be performed. (Total DNA-PKcs stains in a pan-nuclear manner.) As a result, a DNA ligase IV inhibitor (SCR7) was used to impair NHEJ after DNA-PKcs phosphorylation in HFK LXSN cells. Consistent with the hypothesis that impaired NHEJ leads to recruitment of RAD51 when NHEJ cannot be completed, SCR7 increased RAD51/pDNA-PKcs colocalization (Fig 6G-H).

### β-HPV8 E6 abundance correlates with the frequency of DSB repair defects

Immunofluorescence microscopy was used to detect a C-terminal HA tag on the β-HPV8 E6 in HFK β-HPV8 E6 cells (Fig 7A). Analysis using ImageJ software found considerable differences in the intensity of HA staining in these cells (Fig 7B). To determine if cells expressing more β-HPV8 E6 protein were more likely to have DSB repair defects, RAD51 and pDNA-PKcs repair complexes were detected in HFK β-HPV8 E6 cells before and twenty-four hours after DSB induction. HFK β-HPV8 E6 cells that had persistent RAD51 or pDNA-PKcs foci had higher HA-tagged β-HPV8 E6 staining than HFK β-HPV8 E6 cells lacking these unresolved repair complexes (Fig 7C-H, S13A-B Fig.). This was also true for cells with colocalized pDNA-PKcs and RAD51 foci (Fig 7I-K, S13C Fig.). Despite repair defects being more common in cells with elevated HA-tagged β-HPV8 E6, cells with lower HA-tagged β-HPV8 E6 intensity (bottom 50%) had a significantly more persistent/colocalized repair complexes than HFK LXSN cells (S13A-C Fig.). These data support the hypothesis that β-HPV8 E6 drives these repair defects. They also provide evidence that the aberrant repair complexes described here are not the result of an over-expression artefact.

**Fig 7.**
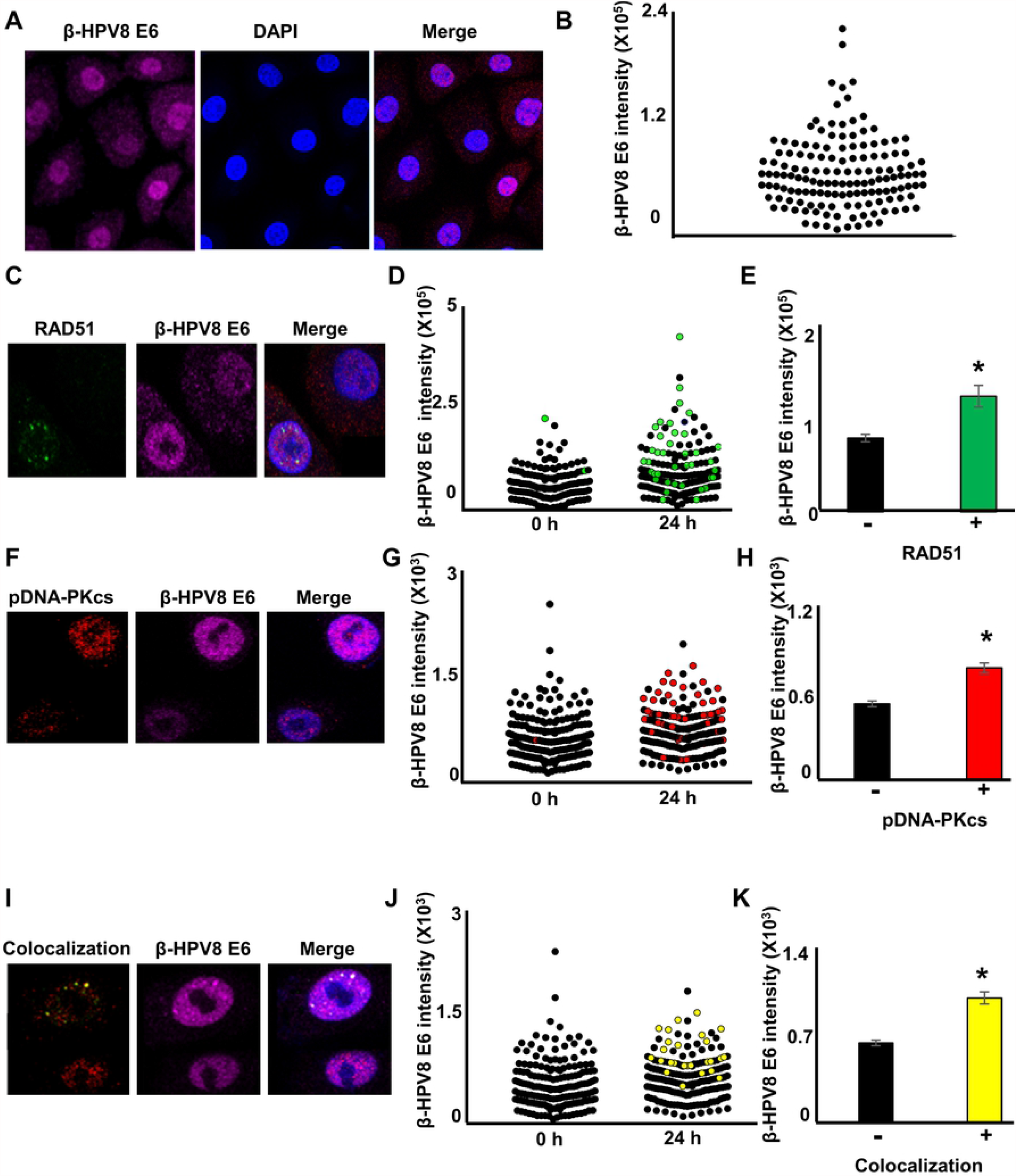
Elevated β-HPV8 E6 expression is associated with DNA repair defects. (A) Representative images of HA-tagged β-HPV8 E6. (B) HA-tagged β-HPV8 E6 intensity in dot plot. (C) Representative images of RAD51 foci and HA-tagged β-HPV8 E6 24-hours after zeocin treatment (10 µg/mL, 10 min). (D) HA-tagged β-HPV8 E6 intensity following zeocin treatment. RAD51 foci positive cells are green. (E) Average β-HPV8 E6 intensity in RAD51 foci negative and RAD51 foci positive cells. (F) Representative images of pDNA-PKcs foci and HA-tagged β-HPV8 E6 24-hours after zeocin treatment. (G) HA-tagged β-HPV8 E6 intensity following zeocin treatment. pDNA-PKcs foci positive cells are red. (H) Average β-HPV8 E6 intensity in pDNA-PKcs foci negative and pDNA-PKcs foci positive cells. (I) Representative images of pDNA-PKcs/RAD51 colocalization and HA-tagged β-HPV8 E6 24-hours after zeocin treatment. (J) HA-tagged β-HPV8 E6 intensity following zeocin treatment. Cells with pDNA-PKcs/RAD51 colocalization are yellow. (K) Average HA-tagged β-HPV8 E6 intensity in cells with or without pDNA-PKcs/RAD51 colocalization. The statistical significance of differences between groups were determined using Student’s t-test. * indicates p < 0.05. For each experiments about 150 cells were imaged over three independent experiments. All microscopy images are 400X magnification.

### Beta-HPV 8 E6 increases genomic instability during DSB repair

The NHEJ and HR pathways are incompatible with each other. HR produces long single-stranded DNA overhangs that facilitate a search for homologous template. In contrast, NHEJ removes single-stranded DNA to produce a substrate for blunt-end ligation. These competing functions suggest that β-HPV8 E6 makes more mutagenic DSB repair in general and more specifically will increase the frequency of deletions. To test this, HFK LXSN and HFK β-HPV8 E6 cells were transfected with vectors that expressed CAS9 endonuclease and sgRNA to induces a DSB just upstream of the CD4 open reading frame [9,59]. A series of overlapping primers targeting the 100 Kb region upstream and downstream of the CAS9 target site were designed (Fig 8A), pooled and used to produce amplicons for next-generation sequencing. The resulting raw reads were trimmed for quality, mapped to the reference sequence and assessed for mutations (SNPs, indels). These data showed a more than ten-fold increase in mutations in HFK β-HPV8 E6 cells compared to HFK LXSN cells (Fig 8B and Fig 9). This includes significantly more replacement, insertion, deletions, multi-nucleotide variation (MNV), and single nucleotide polymorphism (SNP). Consistent with NHEJ and HR both processing the ends of DNA near the DSB, deletions were over twenty-fold more likely in HFK β-HPV8 E6 cells (1579 compared to 75 or 21.1 times more likely). Further, SNPs were over fifteen-fold more likely in HFK β-HPV8 E6 cells (2327 compared to 149 or 15.6 times more likely). These data demonstrate the ability of β-HPV8 E6 to hinder genome maintenance by increasing the frequency of mutations associated with DNA repair.

**Fig 8.**
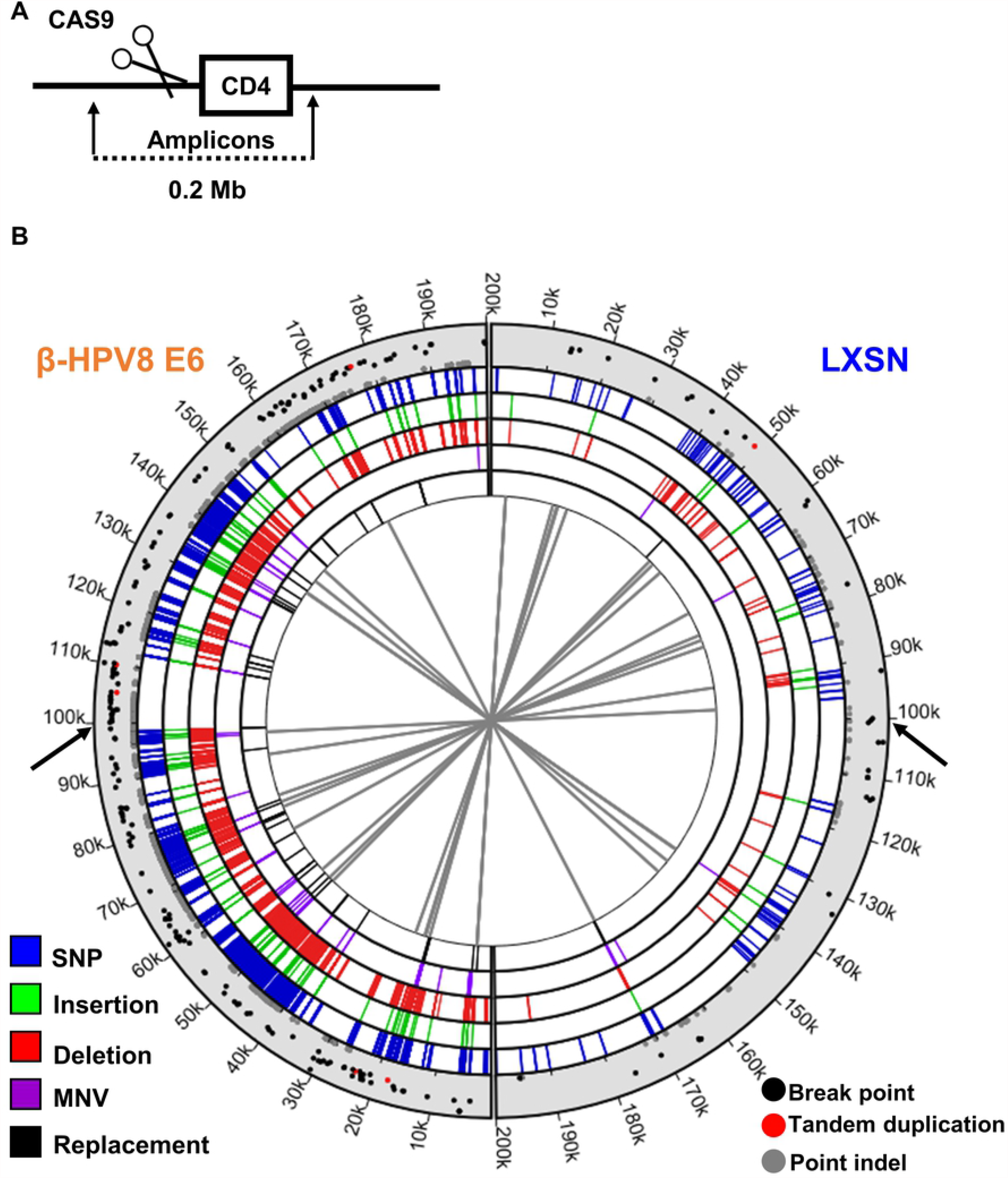
Beta-HPV 8 E6 increases genomic instability during DSB repair. (A) Schematic of the placement of CAS9 induced DSB along the sequenced portion of the genome. (B) Circos plot of DNA mutations in HFK LXSN (right side) and HFK β-HPV8 E6 cells (left side). Black arrows indicate CAS9 cutting sites. The innermost circle displays connections between identical genomic rearrangements. The location of genomic rearrangements colored by types of genomic variations are shown in five concentric circles (blue represents SNP, green represents insertion, red represents deletion, purple represents MNV, and black represents replacement). Scatter plot in the outermost circle displays breakpoints (black), tandam duplications (red), and point indels (grey), in which proximity to the outer edge represents high variant ratio. SNP, single nucleotide polymorphism. MNV, multi-nucleotide variation.

**Fig 9.**
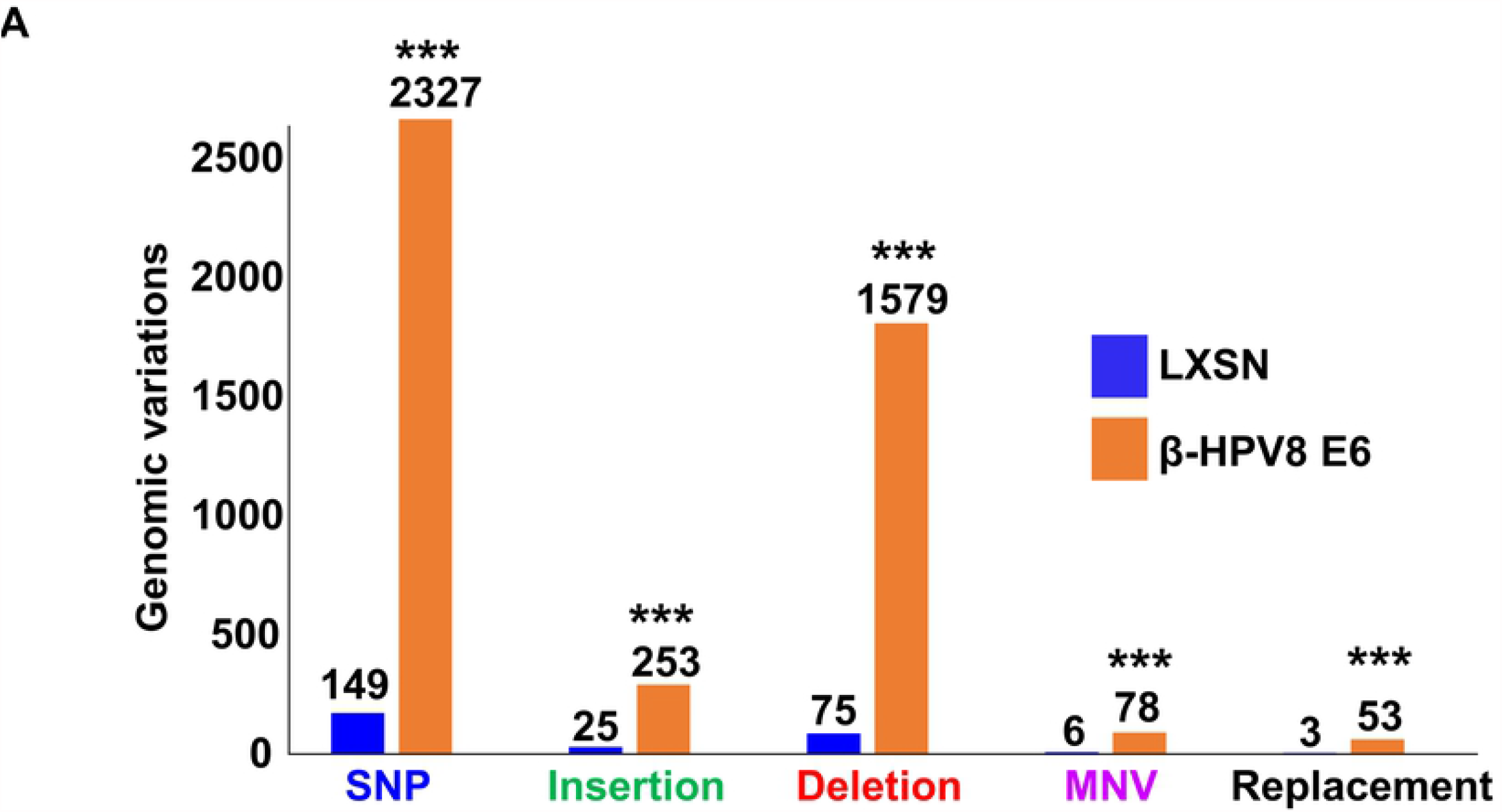
Beta-HPV 8 E6 increases genomic variations during DSB repair. (A) Genomic variations grouped by types of mutational events in HFK LXSN and HFK β-HPV8 E6. Each group of genomic variations and total number of variations were compared between HFK LXSN and HFK β-HPV8 E6. SNP, single nucleotide polymorphism. MNV, multi-nucleotide variation. Statistical differences between cell lines were measured using a Students’ T-test. *** indicates p < 0.001.

## Discussion

We have previously shown that β-HPV8 E6 decreases the efficiency of HR and NHEJ[7,9]. This results in persistent pDNA-PKcs and RAD51 repair complexes and is dependent on β-HPV8 E6 binding and destabilizing p300. Here, we demonstrate that β-HPV8 E6 makes the repair of DSBs significantly more mutagenic. Our work also characterizes the persistent NHEJ and HR repair complexes caused by β-HPV8 E6, demonstrating that HR repair factors (RPA70 and RAD51) are recruited to DSBs when pDNA-PKcs repair complexes do not efficiently repair the lesion. This allows RAD51 repair complexes to form in the G1 phase of the cell cycle. Finally, we provide evidence that p300 plays a critical role in regulating DSB pathway choice, by preventing the initiation of NHEJ and HR at the same lesion.

There are several implications of these data. First, the RPA70/pDNA-PKcs colocalization followed by the formation of RAD51/pDNA-PKcs colocalization suggests cells utilize the HR pathway at lesions where NHEJ fails. In other words, it is possible for repair of a DSB to begin with NHEJ before switching to HR. Supporting this “pathway switch” model, chemical inhibition of NHEJ increased the frequency of RAD51 foci in G1 and the colocalization of RAD51/pDNA-PKcs (Fig 6). As RPA70 foci are an established marker of DNA resection and RAD51 loading requires resected DNA, resection is likely occurring at these lesions [36]. Future studies are needed to characterize DNA end processing during DSB repair in cells expressing β-HPV8 E6. However, the frequency of deletions identified in HFK β-HPV8 E6 cells by our next generation sequencing analysis could be explained by DNA end processing is occurring by both HR and NHEJ pathways. Specifically, we propose that HR creates overhangs and NHEJ removes them, resulting in loss of sequences.

There have been limited reports of RAD51/pDNA-PKcs colocalization. One study found that RAD51/pDNA-PKcs colocalization occurs following a high dosage of UVA radiation [50]. Another study found RAD51/pDNA-PKcs colocalization at common fragile sites in cells exposed to aphidicolin, a DNA polymerase inhibitor [51]. However, neither study characterized these abnormal repair complexes. We show that these colocalization events represent the recruitment of HR factors to sites of stalled or failed NHEJ. Further, p300 plays a critical role in preventing the colocalization of RAD51 and pDNA-PKcs. Since p300 inhibition promoted initiation of HR during G1, CCS1477 may be able to promote gene editing in G1 phase or in non-dividing cells [60,61]. However, this would be contingent on identifying conditions under which the HR pathway could be completed despite p300 inhibition.

Our data are consistent with the proposed role of β-HPV infections in non-melanoma skin cancer development via genome destabilization. We show that β-HPV8 E6 significantly increases the mutational burden of DSBs. We also demonstrate an association between increased β-HPV8 E6 abundance and aberrant DSB repair. This could provide an explanation for why β-HPV infections are more clearly oncogenic in certain populations, where viral loads are elevated, such as people with a rare genetic disease, epidermodysplasia verruciformis, or who are receiving immunosuppressive therapies after organ transplant [1,62,63]. Finally, we have only examined the E6 from β-HPV8. However, the E6 from other members of the β-HPV genus do not destabilize p300, but can immortalize primary cells in combination with expression of β-HPV E7. Thus, continued investigations into the diversity of β-HPV biology are needed to fully evaluate the oncogenic potential of the genus.

## Materials and Methods

### Cell Culture and Reagents

Immortalized human foreskin keratinocytes (HFK), provided by Michael Underbrink (University of Texas Medical Branch), were grown in EpiLife medium (MEPICF500, Gibco), supplemented with 60 µM calcium chloride (MEPICF500, Gibco), human keratinocyte growth supplement (MEPICF500, Gibco), and 1% penicillin-streptomycin (PSL02-6X100ML, Caisson). U2OS and HCT116 cells were maintained in DMEM supplemented with 10% FBS and 1% penicillin-streptomycin. Zeocin (J67140-XF, Alfa Aesar) was used to induce DSBs (10 µg/mL for HFK and 100 µg/mL for U2OS and HCT116 for 10 min). NU7441 (S2638, Selleckchem) was used to inhibit DNA-PKcs phosphorylation (1 µM). CCS1477 (CT-CCS1477, Chemietek) was used to inhibit p300 activity (1 µM). sgRNA/CAS9 plasmids (#JS825, Addgene) were used to generate a DSB for next-generation sequencing.

### Immunofluorescence Microscopy

Cells were seeded onto either 96-well glass-bottom plates and grown overnight. Cells treated with zeocin for specified time and concentration were fixed with 4% formaldehyde. Then, 0.1% Triton-X was used to permeabilize the cells, followed by blocking with 3% bovine serum albumin. Cells were then incubated with the following antibodies: phospho DNA-PKcs S2056 (ab18192, Abcam, 1:200), RAD51 (ab1837, Abcam, 1:200), RPA70 (ab176467, Abcam, 1:200), or HA-tag (#3724, Cell Signalling, 1:100). The cells were washed and stained with the appropriate secondary antibodies: Alexa Fluor 594 (red) goat anti-rabbit (A11012, Thermo Scientific), Alexa Fluor 488 (green) goat anti-mouse (A11001, Thermo Scientific). After washing, the cells were stained with 10 µM DAPI in PBS and visualized with the Zeiss LSM 770 microscope. Images were analysed using the ImageJ techniques previously described [64]. Colocalized foci appear yellow when green and red channels are merged in ImageJ.

### Flow Cytometry

Cells were collected from 10 cm plates, at about 80-90% confluence, by using trypsinization. Cells were washed with cold PBS and fixed with 4% formaldehyde in PBS for 10 min. Then, cells were permeabilize with 0.5% Triton-X for 10 min at room temperature. Cells were stained with anti-RAD51 antibody (ab1837, Abcam, 1:100) and Alexa Fluor 488 goat anti-mouse (A11001, Thermo Scientific,). After washing, cells were resuspended in 200 µL PBS and 30 µM DAPI (4′,6-diamidino-2-phenylindole), and incubated in the dark at room temperature for 15 min. Samples were analysed by a LSRFortessa X20 Flow Cytometer. Flowing software (v2.5.1) was used for data analysis.

### Immunoblotting

After being washed with ice-cold PBS, cells were lysed with RIPA Lysis Buffer (VWRVN653-100ML, VWR Life Science), supplemented with Phosphatase Inhibitor Cocktail 2 (P5726-1ML, Sigma) and Protease Inhibitor Cocktail (B14001, Bimake). The Pierce BCA Protein Assay Kit (89167-794, Thermo Scientific) was used to determine protein concentration. Equal protein lysates were run on Novex 3– 8% Tris-acetate 15 Well Mini Gels (EA03785BOX, Invitrogen) and transferred to Immobilon-P membranes (IPVH00010, Fisher Scientific). Membranes were then probed with the following primary antibodies: GAPDH (sc-47724, Santa Cruz Biotechnologies, 1:1000) and phospho DNA-PKcs S2056 (ab18192, Abcam). P300 (sc-48343, Santa Cruz Biotechnologies), pATM (13050S, Cell signaling), ATM (92356S, Cell Signaling), pATR (58014S, Cell signaling), and ATR (2790S, Cell signaling). After exposure to the matching HRP-conjugated secondary antibody, cells were visualized using SuperSignal West Femto Maximum Sensitivity Substrate (34095, Thermo Scientific).

### SgRNA/CAS9 Transfection

HFK cells were plated in 2 mL of complete growth medium in a 6-well plate. Cells were used at 80-90% confluency. 2 µL of plasmid (#JS825, Addgene) was diluted in 200 µL Xfect transfection reagent (631317, Takara). The mixture was incubated at room temperature for 15 min. The transfection mixture was added to each well drop-wise and incubated for 48 hours at 37 °C. Cells were harvested for DNA extraction and sequencing.

### Next-generation sequencing

Specific primers were designed to cover 0.1 Mb on each side of the Cas9 target site (6689603-6889603 on Chromosome 12) resulting in a total of 42 primer sets each producing a ∼5 Kb overlapping amplicon (S1 Table). Primer sets were pooled based on primer dimerization and annealing temperature compatibility. Genomic DNA was extracted using the MagAttract High Molecular Weight DNA kit (Qiagen) according to the manufacturers’ instructions. The target regions were amplified for each sample using the primer pools coupled with KAPA HiFi Hotstart readymix (KAPA Biosystems) using 20 µM primers as well as a 50/53 °C (Annealing temperature 1/2; Supplementary table 1) and a 5-minute extension time. Primers were removed from PCR amplicons using the Highprep PCR cleanup system (Magbio) as specified by the manufacturer. Libraries were prepared from amplicons with Nextera XT DNA library preparation kit (Illumina) and sequenced on a Nextseq 500 system. A minimum of 100x coverage was targeted for each of the tested samples.

### Sequencing Analyses

Raw reads were trimmed for quality and mapped to the target region in CLC genomics workbench v21.0. Trimmed reads were normalized manually. Normalized reads were assessed for indels and structural variants and normalized for paired read variations in CLC Genomics Workbench v 20.0.4 (Qiagen) using a variant threshold of 5 reads and 100 read coverage (5%). Breakpoints (sites of genomic instability), site-specific variant ratios, insertions, deletions, replacements, inversions and complex (combination of 2 or more genomic changes) were compared between β-HPV8 E6 and the vector control samples.

### Statistical Analysis

All values are represented as mean ± standard error (SE). Statistical differences between groups were measured by using Student’s t-test. p-values in all experiments were considered significant at less than 0.05.

## Acknowledgements

We appreciate KSU-CVM Confocal Core for our immunofluorescence microscopy, Michael Underbrink for providing the TERT-immortalized HFKs, and Jeremy M. Stark for providing plasmids of sgRNA/CAS9. This research was supported by the National Institute of General Medical Sciences (NIGMS) of the National Institutes of Health under award number P20GM130448.

## Data Availability Statement

All relevant data are within the paper and its Supporting Information files.

## Author Contributions

Conceived and designed the experiments: CH NAW RP. Performed the experiments:CH TB RP. Analyzed the data: CH TB RP. Wrote the paper: CH TB NAW RP.

## FIGURES LEGENDS

**S1 Fig**. β-HPV8 E6 allows RAD51 and pDNA-PKcs foci formation to occur in the same cell. (A) Average intensity of HA staining in HFK LXSN and HFK β-HPV8 E6 cells measured by immunofluorescence microscopy over three independent repeats with at least 50 cells per repeat. Statistical significance of the differences between cell lines was determined using Student’s t-test. *** indicates p-value < 1.0 ×10^−50^. (B) Representative images of Rad51 (green) and pDNA-PKcs (red) foci in HFK LXSN and HFK β-HPV8 E6 cells 24-hours after zeocin treatment (10 µg/mL, 10min). (C) Percentage of HFK cells with both Rad51 and pDNA-PKcs foci 24-hours after zeocin treatment. White arrow indicates colocalization. Statistical significance of the differences between cell lines was determined using Student’s t-test. * indicates p-value < 0.05. n=3. All values are represented as mean ± standard error from three independent experiments.

**S2 Fig**. β-HPV8 E6 increases colocalization of RPA70 and RAD51 with pDNA-PKcs in U2OS cells. (A) representative immunoblots showing p300 in U2OS cells. GAPDH is used as a loading control. (B) Representative images of RPA 70 (green) and pDNA-PKcs (red) staining in U2OS LXSN, U2OS β-HPV8 E6, and U2OS β-HPV8 ΔE6 cells following zeocin treatment (100 µg/mL, 10min). (C) Representative images of Rad51 (green) and pDNA-PKcs (red) staining in U2OS LXSN, U2OS β-HPV8 E6, and U2OS β-HPV8 ΔE6 cells following zeocin treatment. White arrow indicates colocalization.

**S3 Fig**. p300 is deleted from HCT116 cells. (A)Immunoblot showing p300 in HCT116 WT and HCT116 p300 knockout (p300 KO) cells. GAPDH is used as a loading control.

**S4 Fig**. p300 inhibitor (CCS1477) decreases phosphorylated ATR (pATR) and phosphorylated ATM (pATM). (A) Representative immunoblots (n=3) showing pATR, ATR, pATM, and ATM expression in cells treated with DMSO or CCS1477 24-hours following zeocin treatment (10 µg/mL, 10min). GAPDH is used as a loading control. This experiment was repeated three times.

**S5 Fig**. p300 inhibitor (CCS1477) increases RPA70/pDNA-PKcs and RAD51/pDNA-PKcs colocalization. (A) Representative images of RPA70 (green) and pDNA-PKcs (red) staining in HFK LXSN cells treated with DMSO or CCS1477 (1 µM) at indicated times following zeocin treatment (10 µg/mL, 10min). (B) Representative images of Rad 51 (green) and pDNA-PKcs (red) staining in HFK LXSN cells treated with DMSO or CCS1477 (1 µM) at indicated times following zeocin treatment (10 µg/mL, 10min). White arrow indicates colocalization.

**S6 Fig**. Negative and positive controls were used to measure RAD51 level in HFK cells using flow cytometry. (A) Representative images of flow cytometry analysis in HFK LXSN and HFK β-HPV8 E6 cells stained only with secondary antibody. (B) Percentage of RAD51 positive HFK LXSN control cells measured by flow cytometry 4-hours following DSB induction by 10 µg/mL zeocin for 10 min. All values are represented as mean ± standard error from three independent experiments. Statistical significance of differences between treatments was determined using Student’s t-test. * indicates p-value < 0.05 (n=3).

**S7 Fig**. β-HPV8 E6 increases RAD51 foci in G1 phase by binding p300. (A) Representative cyclin E positive U2OS LXSN, U2OS β-HPV8 E6, and U2OS β-HPV8 ΔE6 cells stained for Rad51 and cyclin E following zeocin treatment (100 µg/mL, 10min). (B) Representative images of flow cytometry analysis in U2OS LXSN, U2OS β-HPV8 E6, and U2OS β-HPV8 ΔE6 cells stained only with secondary antibody. (C) Representative images of flow cytometry analysis in U2OS LXSN, U2OS β-HPV8 E6, and U2OS β-HPV8 ΔE6 cells stained for DAPI and Rad51 following zeocin treatment. DAPI was used to identify G1 populations (blue area in inset) that were then gated on Rad51 intensity. The gating represents RAD51 positive based off secondary only control.

**S8 Fig**. Negative and positive controls were used to measure RAD51 level in HCT116 cells using flow cytometry. (A) Representative images of flow cytometry analysis in HCT116 p300 WT and HCT116 p300 KO cells stained only with secondary antibody. (B) Percentage of RAD51 positive HCT116 WT cells measured by flow cytometry 24-hours following DSB induction by 100 µg/mL zeocin for 10 min. All values are represented as mean ± standard error from three independent experiments. Statistical significance of differences between groups was determined using Student’s t-test. * indicates p-value < 0.05 (n=3).

**S9 Fig**. p300 inhibition increase RAD51 foci in G1. (A) Representative cyclin E positive HFK LXSN cells treated with DMSO or CCS1477 (1µM) stained for Rad51 (green) and cyclin E (red) following zeocin treatment (10 µg/mL, 10min). (B) Representative images of flow cytometry analysis in HFK LXSN cells treated with DMSO and stained only with secondary antibody. (C) Representative images of flow cytometry analysis in HFK LXSN cells treated with DMSO or CCS1477 (1 µM) and stained for DAPI and Rad51 following zeocin treatment. DAPI was used to identify G1 populations (blue area in inset) that were then gated on Rad51 intensity. The gating represents RAD51 positive based off secondary only control.

**S10 Fig**. Inhibiting DNA-PKcs makes pH2AX persistent following zeocin treatment. (A) Representative immunoblots showing pDNA-PKcs and DNA-PKcs abundance in HFK LXSN cells treated with DMSO or NU7441 following zeocin treatment (10 µg/mL, 10min). β-Actin was used as a loading control. (B) Representative images of HFK LXSN cells treated with NU7441 (1 µM) or DMSO stained for pH2AX S139 (red) following zeocin treatment (10 µg/mL, 10min). (C) Percentage of HFK LXSN cells with pH2AX S139 foci following zeocin treatment. All values are represented as mean ± standard error from three independent experiments. Statistical significance of differences between DMSO and NU7441 treated cells was determined using Student’s t-test. * indicates p-value < 0.05 (n=3).

**S11 Fig**. Negative and positive controls were used to measure RAD51 level in HFK LXSN cells using flow cytometry. (A) Representative images of flow cytometry analysis in HFK LXSN cells treated with DMSO and stained only with secondary antibody. (B) Percentage of RAD51 positive HFK LXSN cells measured by flow cytometry 24-hours following DSB induction by 10 µg/mL zeocin for 10 min. All values are represented as mean ± standard error from three independent experiments. “ns” indicates no significant difference between groups determined using Student’s t-test (n=3).

**S12 Fig**. NU7441 increases RAD51 foci in G1 in U2OS and HCT116 cells. (A) Percentage of cyclin E positive U2OS LXSN cells that have RAD51 foci following zeocin treatment (100 µg/mL, 10min). (B) Percentage of cyclin E positive HCT116 p300 WT cells that have RAD51 foci following zeocin treatment (100 µg/mL, 10min). All values are represented as mean ± standard error from three independent experiments. Statistical significance of differences between cells was determined using Student’s t-test. * indicates p-value < 0.05 (n=3).

**S13 Fig**. Elevated HA-tagged β-HPV8 E6 expression associates with DNA repair defects. HA-tagged β-HPV8 E6 level was defined as “Low β-HPV8 E6” (<median) and “High β-HPV8 E6” (>=median). (A) Percentage of RAD51 foci positive cells in HFK with LXSN control, Low β-HPV8 E6, and High β-HPV8 E6 24-hours after zeocin treatment (10 µg/mL, 10 min). (B) Percentage of pDNA-PKcs foci positive cells in HFK with LXSN control, Low β-HPV8 E6, and High β-HPV8 E6 24-hours after zeocin treatment. (C) Percentage of RAD51/pDNA-PKcs colocalization positive cells in HFK with LXSN control, Low β-HPV8 E6, and High β-HPV8 E6 24-hours after zeocin treatment. All values are represented as mean ± standard error from three independent experiments. Statistical significance in differences of β-HPV8 E6 expressing cells and controls was determined using Student’s t-test. n=3 * indicates p-value < 0.05 (n=3).

